# CK2 Phosphorylation is required for Regulation of Syntaxin 1A activity in Ca^2+^-triggered release in neuroendocrine cells

**DOI:** 10.1101/2020.08.20.258863

**Authors:** Noa Barak-Broner, Dafna Singer-Lahat, Dodo Chikvashvili, Ilana Lotan

**Author notes:** Correspondence should be addressed to: Ilana Lotan, Department of Physiology and Pharmacology, Sackler School of Medicine, Tel Aviv University, Ramat Aviv 69978 Israel.

## Abstract

The polybasic juxtamembrane region (5RK) of the plasma membrane neuronal SNARE, syntaxin1A (Syx), was shown by us to act as a fusion clamp in PC12 cells, making release dependent on stimulation by Ca^2+^. By using a Syx-based FRET probe, we demonstrated that 5RK is absolutely required for a depolarization-induced Ca^+2^-dependent, close-to-open transition (CDO) of Syx that involves the vesicular SNARE synaptobrevin2 and occurs concomitantly with Ca^2+^-triggered release. Here, we investigated the mechanism underlying the 5RK requirement, and identified phosphorylation of Syx at Ser-14 (S14) by protein kinase CK2 as a crucial molecular determinant. Following biochemical verification that both endogenous Syx and CSYS are constitutively S14 phosphorylated in PC12 cells, dynamic FRET analysis of phospho-null and phospho-mimetic mutants of CSYS and the use of a CK2 inhibitor revealed that it is the S14 phosphorylation that confers the 5RK requirement. Concomitant amperometric analysis of catecholamine release revealed that the phospho-null mutants do not support release, spontaneous and evoked. Collectively, these results identify a functionally important CK2 phosphorylation site in Syx that is required for 5RK-regulation of CDO and for concomitant Ca^2+^-triggered release.

**Summary statement:** Many phospho-proteins participate in vesicle exocytosis. We show that a recently identified structural transition of syntaxin1A that accompanies Ca^2+^-regulated exocytosis in neuroendocrine cells is controlled by CK2 phosphorylation of syntaxin1A.

## Introduction

Ca^2+^-triggered exocytosis is a fundamental cellular process mediated by the SNARE proteins, syntaxin 1A (Syx), synaptobrevin2 (Syb2), and SNAP-25 (Rizo and Xu, 2015; Rothman, 1996). The plasma membrane (PM) SNARE, Syx, exists in two alternative conformations: a ‘closed’ conformation in which the SNARE motif is blocked by the Habc-domain (Fernandez et al., 1998), and an ‘open’ conformation in which the SNARE motif is exposed. In the ‘open’ conformation Syx can participate in the formation of SNARE complexes (Dulubova et al., 2007).

The very N terminal end of the Syx protein contains a highly conserved domain of 27 amino acids (N-peptide) (Rickman et al., 2007), whose binding to the auxiliary SM protein Munc18-1 is essential for fusion (Deak et al., 2009; Khvotchev et al., 2007; Rathore et al., 2010). In addition, the N-peptide, harbors a specific region of interest at Ser-14 (S14), which is the sole target in Syx for phosphorylation by casein kinase 2 (CK2) (Dubois et al., 2002; Rickman and Duncan, 2010), a pleiotropic constitutively active serine/threonine protein kinase with particularly high levels in brain (Cozza et al., 2012). A number of observations over the past few years have suggested a potential role for CK2 phosphorylation of S14 in Syx in the direct modulation of exocytosis; i) S14 phosphorylation increased through development in parallel with synapse development and maturation in rat brains (Foletti et al., 2000); ii) S14 phosphorylation enhanced the interaction between Syx and synaptotagmin1 (Syt1) *in vitro* (Risinger and Bennett, 1999); iii) modification of S14 was found to play a role in a specific mode of the Syx-Munc18-1 interactions, affecting vesicle immobilization and exocytosis in PC12 cells (Rickman and Duncan, 2010).

Another structural determinant that is important for the activity of Syx in exocytosis is the polybasic stretch of 5 lysines and arginines (5RK) located at the juxtamembrane linker region that connects the SNARE motif and the transmembrane domain (TMD). This 5RK region was recognized as a lipid binding domain that plays a critical role in determining the energetics of SNARE-mediated fusion events (Lam et al., 2008) and is essential for lipid mixing and for the transition from hemifusion to full fusion in liposomes (Hernandez et al., 2012). In PC12 cells, electrostatic interactions of 5RK with PI(4,5)P2 (PIP2) are thought to mediate the clustering and segregation of Syx into distinct microdomains where synaptic vesicles undergo exocytosis (James et al., 2009; Murray and Tamm, 2009; van den Bogaart et al., 2011), possibly by serving as molecular beacons for vesicle docking by a Syt1 interaction with 5RK through PIP2 (Honigmann et al., 2013). Similarly, electrostatic interactions between 5RK and PI(3,4,5)P3 have also been suggested to facilitate Syx clustering (Khuong et al., 2013).

Previously, in an attempt to gain insights into the domain-specific-secretion-related functioning of Syx, we constructed a Syx-based fluorescence resonance energy transfer (FRET) probe, which allowed us to resolve a depolarization-induced Ca^2+^-dependent close-to-open transition (CDO) of Syx that results from Ca^2+^ entry through voltage gated Ca^2+^ channels (Greitzer-Antes et al., 2013). CDO was accompanied by Ca^2+^-triggered release, both in PC12 cells (Greitzer-Antes et al., 2013) and hippocampal neurons (Vertkin et al., 2015). Notably, CDO was absolutely dependent on intact 5RK (Greitzer-Antes et al., 2013) and on plasma membrane PIP2 (Singer-Lahat et al., 2018). Further, our results suggested that the role of 5RK in exocytosis may be more central than has been previously demonstrated (Khuong et al., 2013; Lam et al., 2008; Murray and Tamm, 2009). Indeed, amperometric analysis in PC12 cells revealed that charge neutralization of 5RK promoted spontaneous release and inhibited Ca^2+^-triggered release, indicating that 5RK acts as a fusion clamp, making release dependent on stimulation by Ca^2+^ (Singer-Lahat et al., 2018)

This current report identifies phosphorylated S14 as a significant molecular determinant in Syx that enables CDO to occur within a vesicular setting in which it is absolutely dependent on the presence of intact 5RK, supporting catecholamine release. We propose a model that describes the roles of phosphorylated S14 and 5RK and their interplay in exocytosis.

## Results

Previously, we used a Syx-based FRET probe, CSYS (Fig. 1A), which reports Ca^2+^-dependent close-to-open transition of Syx in response to high K^+^ (hK) depolarization (termed: Ca^2+^-dependent opening; CDO), to reveal that the polybasic juxtamembrane 5RK stretch of Syx is essential for CDO and consequently for Ca^2+^-triggered release (Singer-Lahat et al., 2018). Here, acknowledging the potential relevance of CK2-induced S14 phosphorylation of Syx to exocytosis (‘*Introduction’*), we used dynamic FRET and amperometry analyses to explore the possibility that S14 phosphorylation is involved in the molecular/cellular events that endow 5RK its central role.

**Figure 1.**
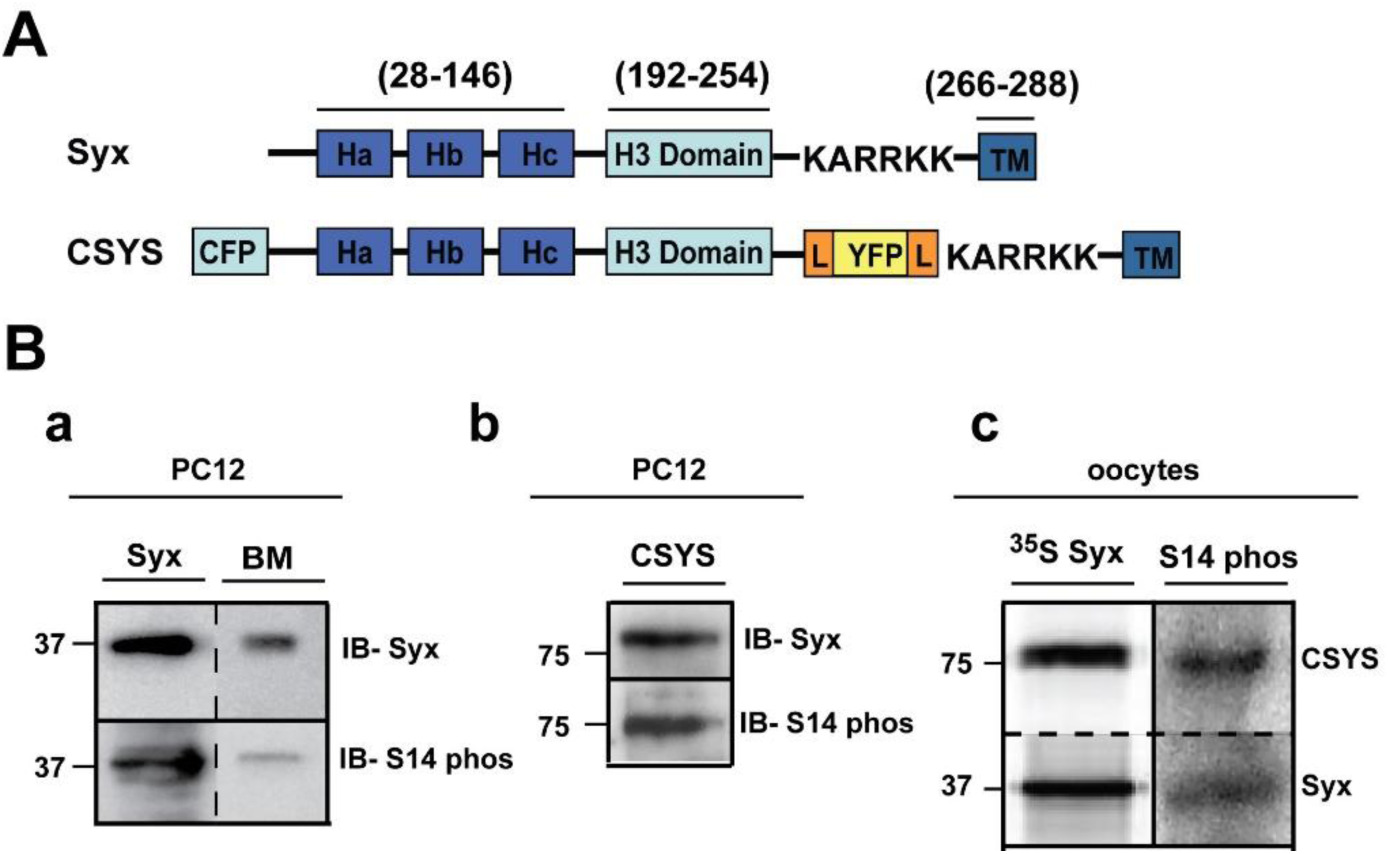
Syx and CSYS are targets for CK*2* phosphorylation. (**A**) Domain structure of Syx and the CSYS FRET probe. (**B**) CSYS is a target for CK2 phosphorylation similarly to native Syx. **(a)** Native PC12 cells were immunoprecipitated with anti-Syx antibody and immunoblotted either with anti-Syx antibody (*IB-Syx*; upper panel) or with Syx S14 phospho-specific antibody (*IB-S14 phos*; lower panel). Brain membranes (BM) were loaded as a control. (**b)** Following immunoprecipitation with anti-YFP antibody, proteins of PC12 cells transfected with CSYS were subjected to Western blot analysis with either anti-Syx antibody (*IB-Syx*; upper panel) or with Syx S14 phospho-specific antibody (*IB-S14 phos*; lower panel). Molecular markers are shown on the left. **(c)** *Xenopus* oocytes expressing CSYS and Syx proteins were immunoprecipitated with anti-Syx antibody (^*35*^*S Syx*; left panel) and immunoblotted with Syx S14 phospho-specific antibody (*IB-S14 phos*; right panel).

### S14 phosphorylation of CSYS is the molecular determinant that makes 5RK essential for CDO

First, using S14 phospho-specific antibody in a Western blot analysis, we established that in the PC12 cells used in our experiments endogenous Syx is S14 phosphorylated (Fig. 1Ba). Also, CSYS, exogenously expressed in the same cell, is S14 phosphorylated (Fig. 1Bb). The extent of phosphorylation of CSYS is similar to that of Syx, as determined in the heterologous expression system of *Xenopus* oocytes (Fig. 1Bc).

Next, we set to explore the possible involvement of S14 phosphorylation in CDO using the S14 phospho-null mutant CSYS;S14A (Fig. 2A; S14 was mutated to alanine to prevent phosphorylation by CK2 (Rickman and Duncan, 2010)). We have previously defined CDO in CSYS expressing cells, using dynamic FRET analysis, as the reduction in the F_YFP_/F_CFP_ ratio (FRET ratio), reporting a close-to open transition, that occurs in response to hK depolarization and is blocked by the voltage-gated Ca^2+^-channel blocker, cadmium (Cd) (Greitzer-Antes et al., 2013). Here, monitoring the FRET ratio reduction in response to hK depolarization in CSYS;S14A expressing cells, we found that it was similar to that monitored in parallel in CSYS expressing cells (Fig. 2Ba) and consisted of a fraction (CDO) that was blocked by 200 μM Cd (Fig. 2Bb; F(79,6083) = 15.116, p < 0.0001, ANOVA with repeated measures). This analysis suggested that CSYS;S14A undergoes CDO, similarly to CSYS.

**Figure 2.**
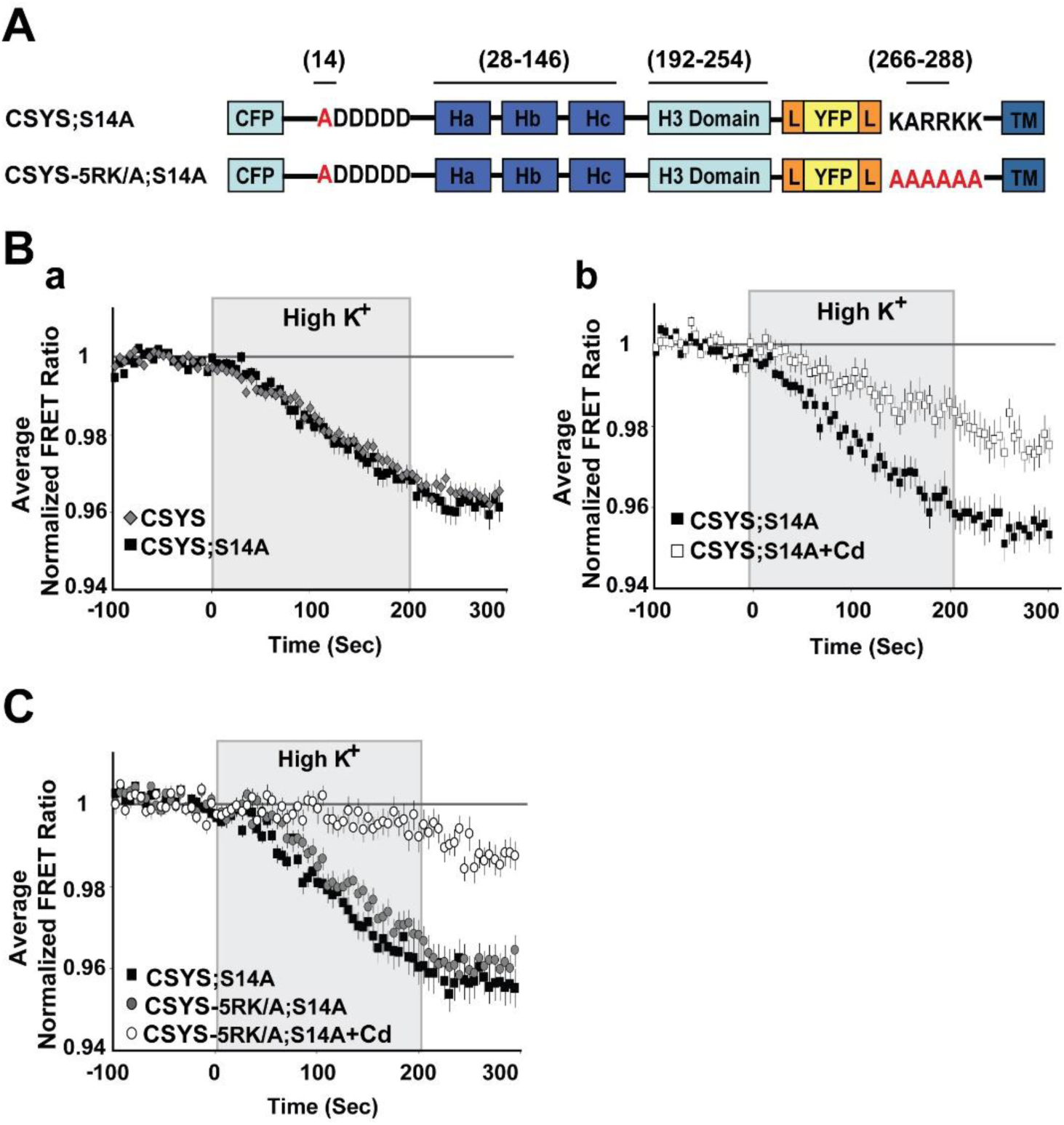
The phospho-null mutant undergoes CDO that does not require intact 5RK. **(A)** Domain structure of the CSYS mutants; CSYS;S14A and CSYS-5RK/A;S14A. (**B)** The phospho-null mutant CSYS;S14A undergoes CDO. (**a)** The reduction in the FRET ratio of the phospho-null mutant CSYS;S14A in response to hK depolarization is similar to that of CSYS (black squares, *n = 30* cells, and grey diamonds *n = 32* cells, respectively; 3 experiments). **(b)** Part of the FRET ratio reduction of CSYS;S14A is blocked by Cd and represents CDO. Addition of 200 µM Cd to both control and hK solutions resulted in a smaller reduction in the FRET ratio (white squares, *n* = 22 cells, and black squares, *n* = 26 cells; 2 experiments; F(79,6083) = 15.166, p < 0.0001, ANOVA with repeated measures). **(C)** CDO in the phospho-null mutant does not require intact 5RK. Despite the 5RK/A mutation, the reduction in FRET ratio of CSYS-5RK/A;S14A in response to hK depolarization was similar to that of CSYS;S14A (grey circles, *n = 39*, and black squares, *n = 44*, respectively; 5 experiments) and was smaller in the presence of Cd (white circles *n = 40*, 5 experiments; F(79,3397) = 5.411, p < 0.0001, ANOVA with repeated measures).

Absolute requirement of intact 5RK for CDO was previously demonstrated with the 5RK charge neutralization mutant, CSYS-5RK/A (the five basic residues replaced with alanines), in which the FRET ratio reduction in response to hK depolarization was smaller than that in CSYS and resistant to Cd block; namely the neutralization of 5RK completely abolished CDO (Greitzer-Antes et al., 2013). To explore the possible involvement of S14 phosphorylation in the requirement of 5RK for CDO, we generated a phospho-null version of CSYS-5RK/A, CSYS-5RK/A;S14A (Fig. 2A), and subjected cells expressing the mutant to dynamic FRET analysis, in parallel to CSYS;S14A. Strikingly, the FRET ratio reduction of CSYS-5RK/A;S14A in response to hK depolarization was similar to that of CSYS;S14A and consisted of CDO, a fraction that was blocked by Cd (Fig. 2C; F(79,3397) = 5.411, p < 0.0001, ANOVA with repeated measures). Thus, unlike the case with CSYS, in the phospho-null mutant intact 5RK is not required for CDO, strongly suggesting that S14 phosphorylation is the molecular determinant responsible for the requirement of 5RK for CDO.

To substantiate this notion, we repeated the same experiments, but with the phospho-mimetic mutation S14E (serine 14 replaced by glutamate). In CSYS;S14E (Fig. 3A) the FRET ratio reduction in response to hK depolarization was similar to that of CSYS (Fig. 3Ba), as was the case with CSYS;S14A. However, the neutralization of 5RK in CSYS-5RK/A;S14E (Fig. 3A), in contrast to CSYS-5RK/A;S14A, but like in CSYS-5RK /A, caused a significantly smaller FRET ratio reduction (Fig. 3Bb; F(79,2844) = 4.331, p < 0.0001, ANOVA with repeated measures). This suggested that CDO was eliminated in the phospho-mimetic mutant in the absence of intact 5RK, in accord with the notion that it is the S14 phosphorylation that makes 5RK a prerequisite for CDO.

**Figure 3.**
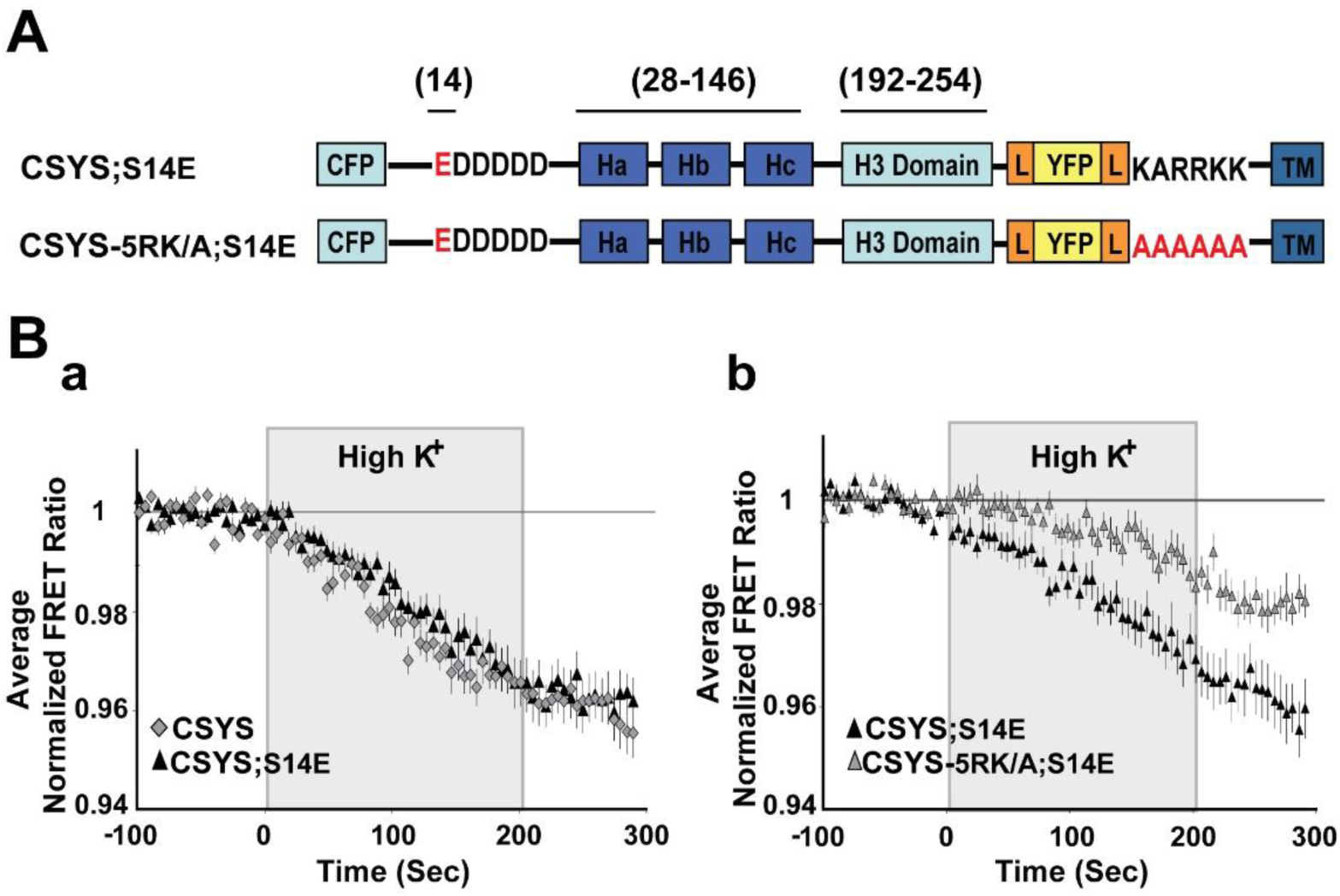
The phospho-mimetic mutant undergoes CDO that requires intact 5RK. **(A)** Domain structure of the phospho-mimetic CSYS mutants, CSYS;S14E and CSYS-5RK/A;S14E. **(B)** The reduction in the FRET ratio in response to hK depolarization of CSYS;S14E was similar to that of CSYS (**a**; black triangles, *n = 18*, and grey diamonds *n = 20*, respectively; 2 experiments) and included CDO that requires intact 5RK, similarly to CSYS, as it was smaller upon insertion of the 5RK/A mutation (**b**; black triangles *n = 18*, and grey triangles, *n = 20*; 2 experiments; F(79,2844) = 4.331, p < 0.0001, ANOVA with repeated measures).

As further validation, we chose to test whether elimination of S14 phosphorylation by endogenous CK2 would also eliminate the requirement of 5RK for CDO. We used two approaches to impair endogenous CK2 phosphorylation. First, we targeted the polyacidic N-terminus stretch of Syx (5D), which is a highly-conserved recognition site for CK2 (Rickman and Duncan, 2010). Dynamic FRET analysis of CSYS-5D/A, with all 5 aspartates neutralized to alanines (Fig. 4A), showed that the FRET ratio reduction in response to hK depolarization was similar to that of CSYS (Fig. 4B). However, that of CSYS-5RK/A;5D/A was significantly larger than that of CSYS-5RK/A and was sensitive to Cd blockade (Fig. 4C; F(79,5214) = 5.576 and F(79,5688) = 8.959; respectively, p < 0.0001, ANOVA with repeated measures), indicating that the 5RK/A mutation did not eliminate CDO in the 5D/A mutant. This data indicate that prevention of S14 phosphorylation *in vivo* by abrogating the CK2 recognition site eliminates the requirement of intact 5RK for CDO, essentially mimicking the effect obtained with the S14A mutation.

**Figure 4.**
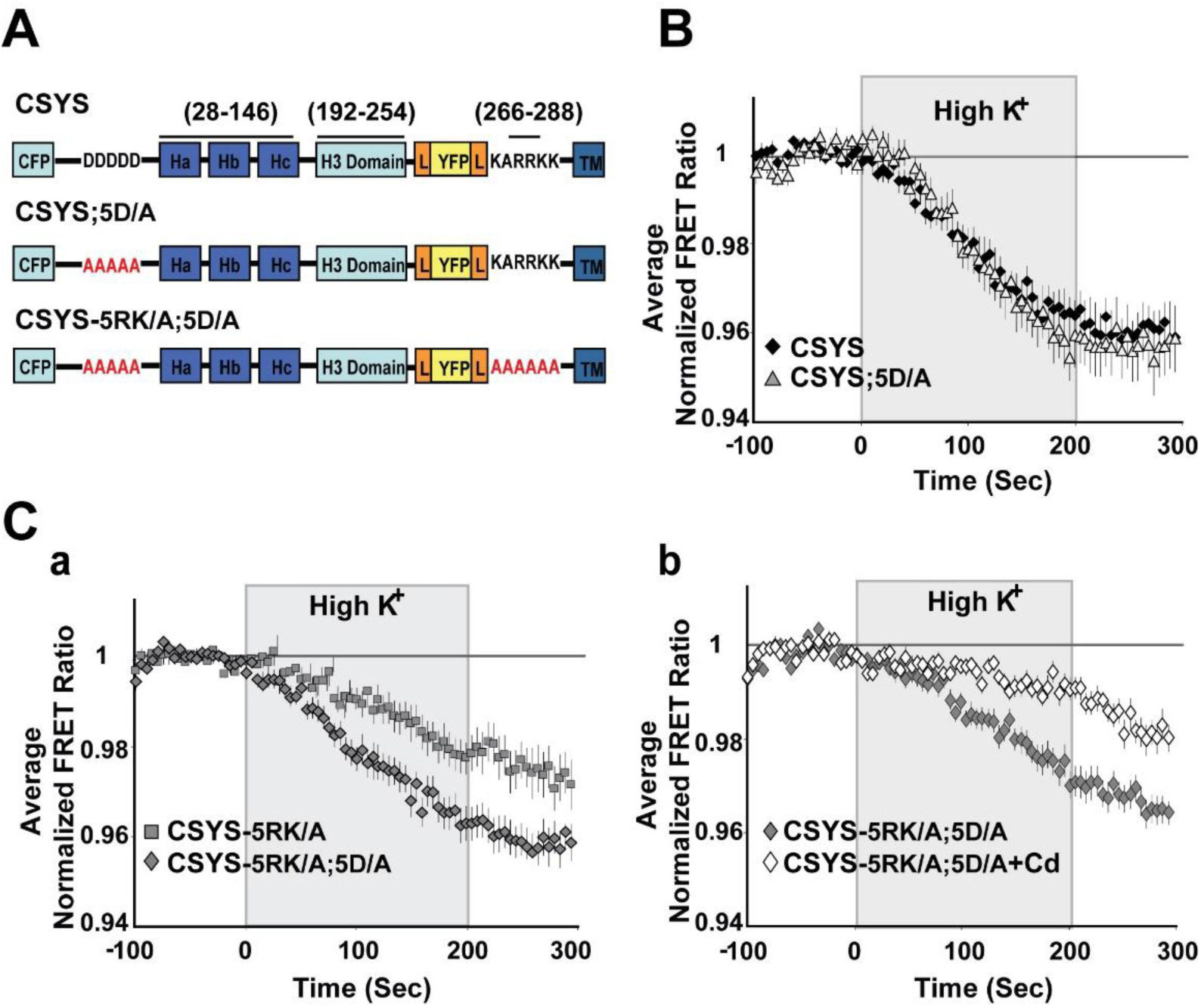
Neutralization of the N-terminal CK2 recognition site in CSYS mimics the effect captured by the phospho-null mutant, eliminating the requirement of 5RK for CDO. **(A)** Domain structures of the CSYS mutants, CSYS;5D/A and CSYS-5RK/A;5D/A. **(B)** The FRET ratio reduction in response to hK depolarization of CSYS;5D/A was similar to that of CSYS (grey triangles, n = 38 and black diamonds, n = 65, respectively; 7 experiments). **(C)** The FRET ratio reduction of CSYS-5RK/A;5D/A was larger than that of CSYS-5RK/A (**a;** grey diamonds, n = 35, and grey squares n = 34, respectively; 7 experiments; F(79,5214) = 5.576, p < 0.0001, ANOVA with repeated measures) and was smaller in the presence of Cd, representing CDO that does not require intact 5RK (**b;** white diamond, n = 39, and grey diamonds, n = 42, respectively; 3 experiments; F(79,5688) = 8.959, p < 0.0001, ANOVA with repeated measures).

The second approach to impair the *in vivo* S14 phosphorylation involved inhibition of CK2 function by tetrabromo-2-benzotriazole (TBB), a highly selective ATP/GTP competitive inhibitor of CK2 (Sarno et al., 2001). Western blot analysis verified that 20 min incubation of PC12 cells with 20µM TBB, inhibited efficiently phosphorylation of both endogenous Syx and over-expressed CSYS-5RK/A (Fig. 5A). Using this approach, we challenged CSYS-5RK/A, which normally does not undergo CDO (Greitzer-Antes et al., 2013), with the expectation that the prevention of its *in vivo* phosphorylation by TBB would remove the requirement of 5RK for CDO. Remarkably, incubation with TBB increased the FRET ratio reduction in response to hK depolarization of CSYS-5RK/A, and the effect was Cd-sensitive (Fig 5B; F(79,3239) = 3.239 and F(79,5214) = 3.410; respectively, p < 0.0001, ANOVA with repeated measures), indicating that CDO was enabled by the impairment of S14 phosphorylation, as expected.

**Figure 5.**
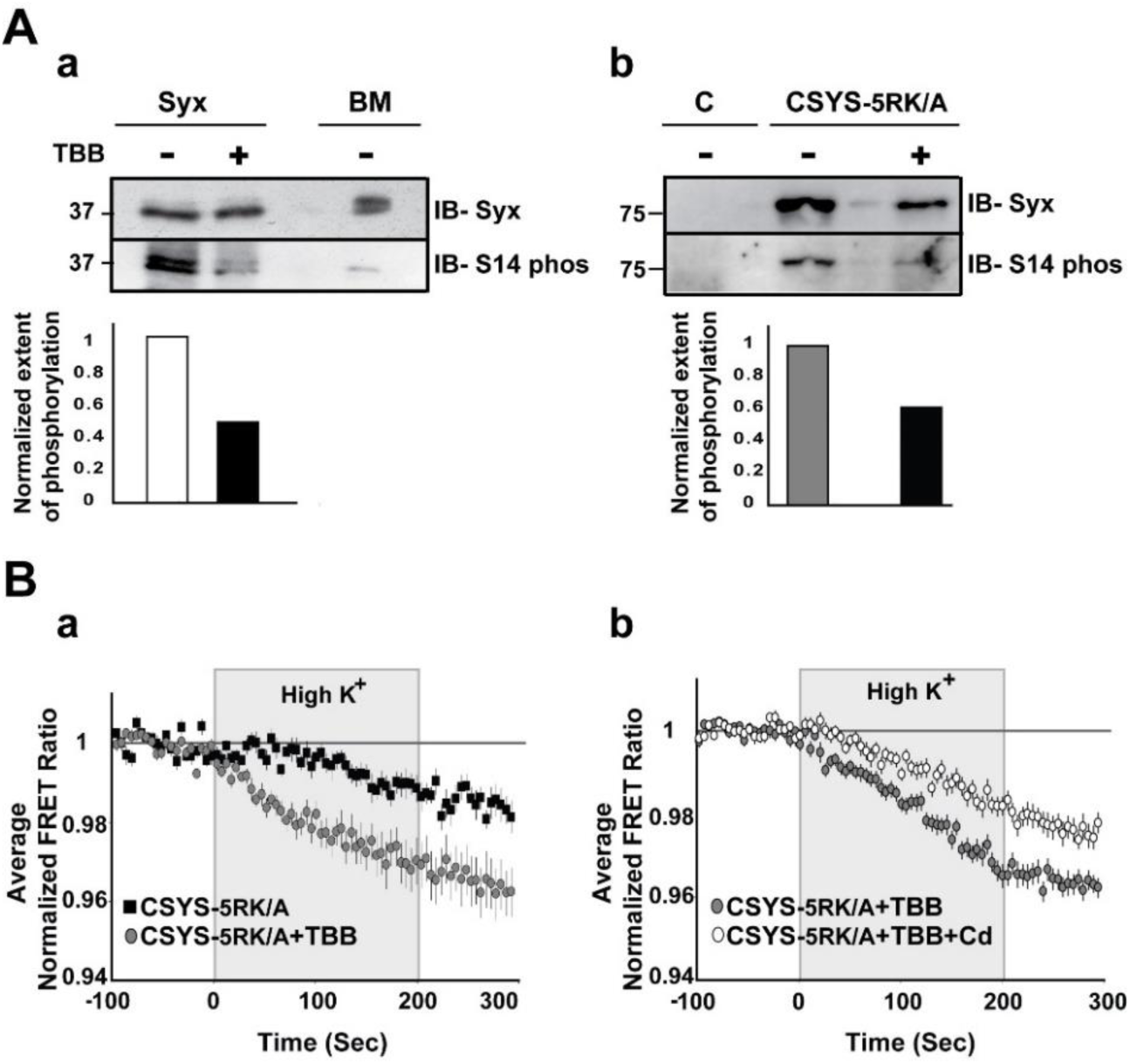
Inhibition of *in vivo* S14 phosphorylation eliminates the requirement of 5RK for CDO. **(A**) **S14 phosphorylation is inhibited by TBB. (a)** S14 phosphorylation of endogenous Syx in PC12 cells is reduced by TBB. Native Syx was immunoprecipitated with anti-Syx antibody and immunoblotted with either anti-Syx antibody (*IB-Syx*; upper panel) or with Syx S14 phospho-specific antibody (*IB-S14 phos*; middle panel). The extent of phosphorylation (ratio between the corresponding upper and middle intensities) was reduced by TBB (lower panel). Native brain membranes (BM) were loaded as a control. **(b)** S14 phosphorylation of CSYS-5RK/A expressed in PC12 cells is reduced by TBB. CSYS-5RK/A was immunoprecipitated from transfected PC12 cells with anti-YFP antibody and immunoblotted with either anti-Syx antibody (*IB-Syx*; upper panel) or with Syx S14 phospho-specific antibody (*IB-S14 phos;* middle panel). In the presence of TBB the extent of S14 phosphorylation was reduced (lower panel). Molecular weight markers are shown on the left. Native cells were used for control (c). **(B) De-phosphorylation by TBB rescues CDO in CSYS-5RK/A. (a)** Addition of TBB increased the FRET ratio reduction of cells expressing CSYS-5RK/A in response to hK depolarization (grey circles, n = 25, and black squares, n = 19; 2 experiments; F(79,3239) = 3.454, p < 0.0001, ANOVA with repeated measures) **(b)** In the presence of TBB the FRET ratio reduction was smaller upon addition of Cd (white circles n = 34, and grey circles n = 34, respectively; 3 experiments; F(79,5214) = 3.410, p < 0.0001, ANOVA with repeated measures).

Taken together, the results derived using the phospho-null (Fig. 2) and phospho-mimetic (Fig. 3) CSYS mutants with the results of impaired *in vivo* phosphorylation (Figs. 4, 5), strongly implicate CK2-induced S14 phosphorylation as a regulatory mechanism that imposes absolute dependence on intact 5RK for CDO.

### The requirement of PIP2 for CDO is not linked to S14 phosphorylation

Our previous work suggested that CDO is absolutely dependent on PIP2 levels in the PM of PC12 cells (Singer-Lahat et al., 2018). Here, having established the importance of S14 phosphorylation for the requirement of 5RK, we next asked whether phosphorylation could also be linked to the PIP2 requirement. Specifically, we asked whether CDO in the phospho-null mutants, CSYS;S14A and CSYS-5RK/A;S14A, will be eliminated upon hydrolysis of PIP2 by co-expression of the strongly Ca^2+^-activated phospholipase, Cη2 (PLCη2) (Kabachinski et al., 2016). Dynamic FRET analysis in both mutants revealed that the FRET ratio reductions in response to hK depolarization were significantly smaller in the presence of the mRFP-tagged phospholipase (Fig. 6A; F(79,6794) = 8.167 and B; F(79,2686) = 5.343, p < 0.0001, ANOVA with repeated measures); that of CSYS-5RK/A;S14A was verified not to be sensitive to Cd Blockade (Fig. 6C), indicating that CDO was eliminated by the phospholipase. Of note, we previously showed that the effect of PLCη2 captured in our experimental system could be fully attributed to its PIP2 hydrolysis function, and not to the generation of secondary messengers (Singer-Lahat et al., 2018). Taken together, in contrast to the requirement of 5RK, the requirement of PIP2 was preserved in the phospho-null mutant, suggesting that it is not conferred by S14 phosphorylation, but rather may reflect an inherent characteristic of CDO.

**Figure 6:**
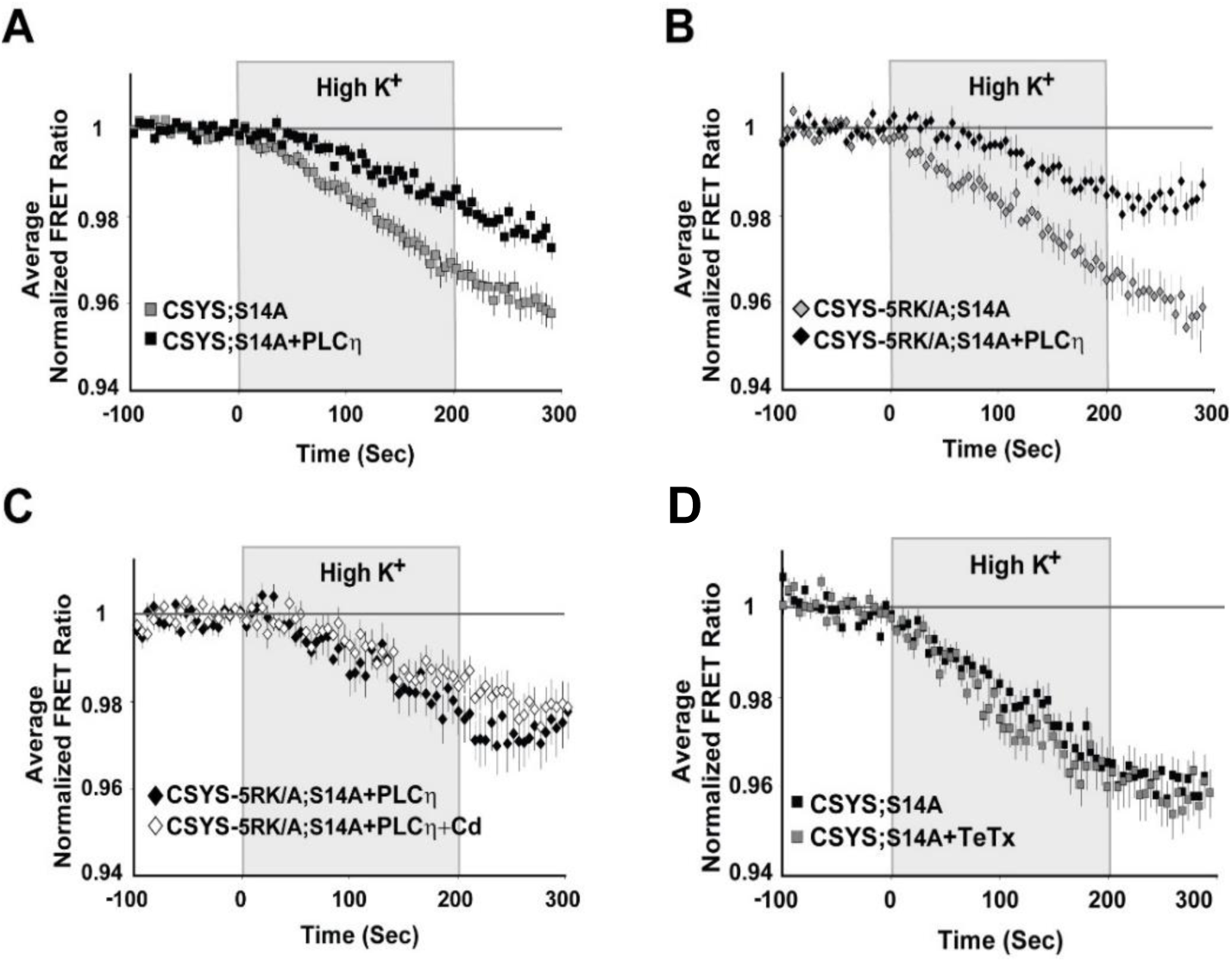
(A-C) A requirement for PIP2 is an intrinsic feature of CDO and is independent of S14 phosphorylation. Upon PIP2 hydrolysis by co-expressed PLCη2, the FRET ratio reduction in response to hK depolarization was reduced in cells expressing each of the phospho-null mutants, CSYS;S14A (**A**; grey squares, n = 50, and black squares, n = 42, respectively; 5 experiments; F(79,6794) = 8.167, p < 0.0001, ANOVA with repeated measures) and CSYS-5RK/A;S14A (**B;** black diamonds, n = 16 and grey diamonds, n = 20, respectively; 2 experiments; F(79,2686) = 5.343, p < 0.0001, ANOVA with repeated measures). The FRET ratio reduction of CSYS-5RK/A;S14A in the presence of PLCη2 was insensitive to the addition of Cd, indicating that PLCη2 inhibited CDO (**C**; white diamonds, n = 21 and black diamonds, n = 23, respectively; 2 experiments). **(D) The conformational transition of the CSYS phospho-null mutant does not involve Syb2**. Syb2 cleavage by TeTx-LC did not have any effect on the FRET ratio reduction in response to hK depolarization of CSYS;S14A (black squares, n = 23 and grey squares, n = 20, respectively; 3 experiments).

### S14 phosphorylation is required for CDO to occur within a vesicular context

Having recognized the significance of S14 phosphorylation for CDO, the next step was to understand its relevance to vesicle release. Previously, we have shown that CSYS, which is constitutively S14 phosphorylated *in vivo* (see Fig. 1B), undergoes CDO that occurs in a vesicular context, as it was eliminated following 4 hours incubation with 30 nM light chain of tetanus toxin (TeTx LC), which cleaves the vesicular SNARE Syb2 (Singer-Lahat et al., 2018). Here, we examined whether the CDO of the phospho-null mutant, CSYS;S14A, also involves Syb2. Strikingly, in contrast to CSYS, there was no effect of TeTx LC on the FRET ratio reduction in response to hK in the phospho-null mutant (Fig 6D), indicating that its opening, including CDO, does not occur in a vesicular context.

### S14 phosphorylation is required for Ca^2+^-triggered release

Previously, an amperometric analysis of catecholamine release showed that 5RK plays a central role in clamping spontaneous release and promoting Ca^2+^-triggered release in PC12 cells, as the 5RK/A mutation enhanced spontaneous release and inhibited Ca^2+^-triggered release (Singer-Lahat et al., 2018). To assess the relevance of S14 phosphorylation to secretion directly, we used carbon fiber amperometry to analyze the secretion phenotype of the phospho-null mutant CSYS;S14A and compared it with the previously characterized (Singer-Lahat et al., 2018) phenotype of CSYS. In order to reduce potentially confounding effects from endogenous Syx, we used cells transfected with the light chain of BoNT-C1α51 (BoNT-C1), which spares SNAP-25 cleavage and cleaves only endogenous Syx (Wang et al., 2011), precluding Syx from mediating membrane fusion (Schiavo et al., 1995). We have previously shown that CSYS(R), a CSYS mutant bearing the K253I mutation, which confers resistance to BoNT-C1 (Lam et al., 2008), is indeed resistant to the co-expressed toxin and retains its PM expression in PC12 cells. Importantly, CSYS(R) was shown to rescue the inhibition of exocytosis by the toxin, in both PC12 cells (Greitzer-Antes et al., 2013) and in hippocampal neurons (Vertkin et al., 2015). Accordingly, we generated here the toxin resistant version of the phospho-null mutant, CSYS;S14A(R), and verified that it is resistant to the toxin and is expressed on the PM of cells co-expressing the toxin (Fig. 7A). Comparative amperometric measurements, 1 min before (to detect spontaneous events) and 1 min after a 10 sec application of hK solution (to detect evoked events), performed in cells co-expressing the toxin with CSYS;S14A(R) or CSYS(R) in the same experiments, revealed that cells expressing either construct had few spontaneous release events (Fig. 7B). Notably, in contrast to CSYS(R), which exhibited evoked release in response to hK depolarization, there was no significant change in the release pattern of CSYS;S14A(R) expressing cells in response to hK depolarization, neither in number of spikes or total charge release (Fig. 7B,C), indicating that the phospho-null mutation eliminated evoked release. Furthermore, cells expressing the toxin resistant version of the double mutant CSYS-5RKA;S14A(R) did not display enhanced spontaneous release (Fig. 7D), as was observed in cells expressing CSYS-5RK/A(R) (Singer-Lahat et al., 2018); namely, the phospho-null mutation eliminated also the enhanced spontaneous release conferred by the 5RK/A mutation.

**Figure 7.**
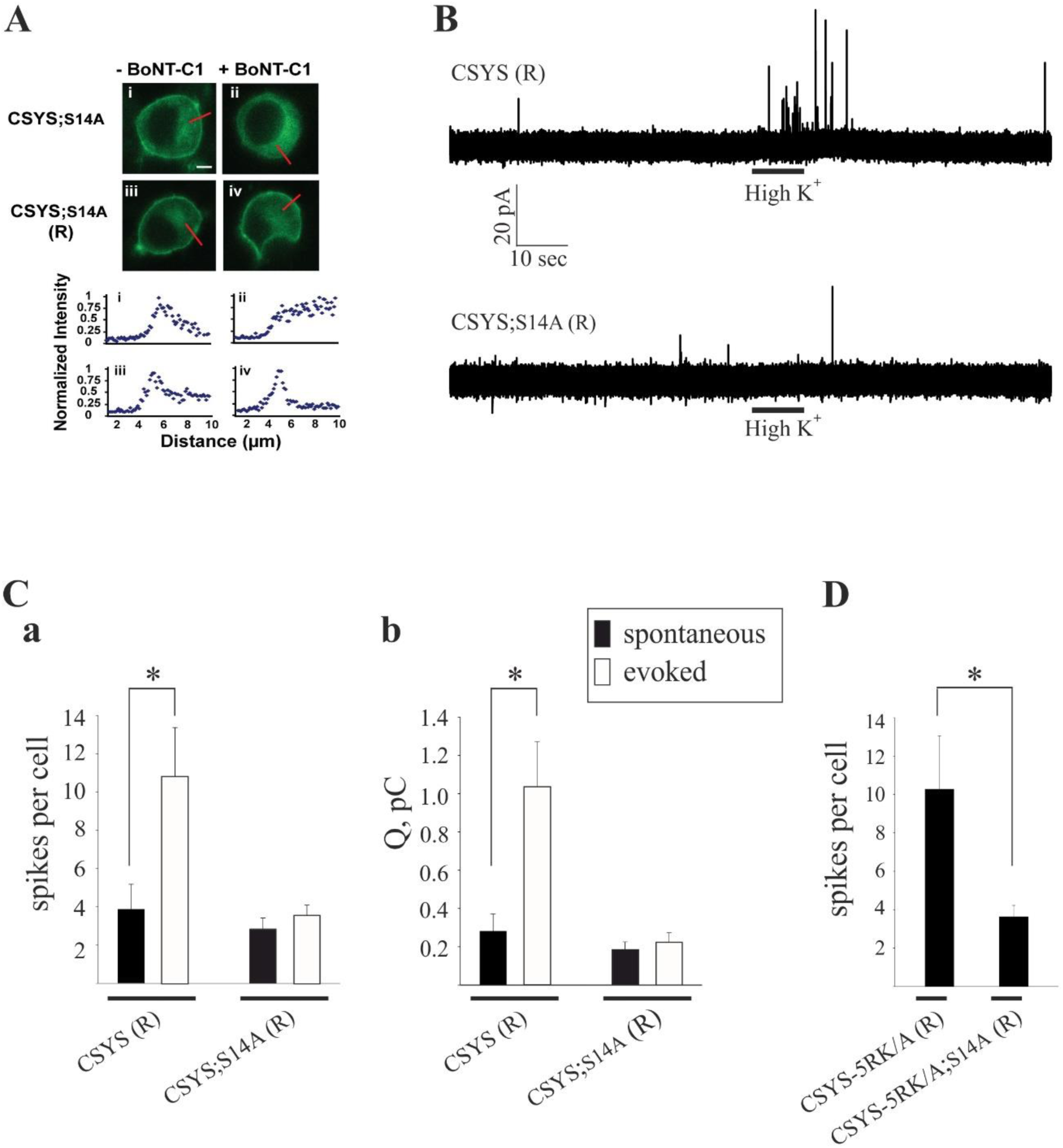
The phospho-null mutation abolishes Ca^2+^-triggered release. **(A**) CSYS;S14A (R), bearing the K253I mutation, is located in the PM region of PC12 cells and is resistant to cleavage by BoNT-C1. *Upper panel*: confocal images of PC12 cells demonstrating that CSYS;S14A (R) (iii) retained its PM distribution upon BoNT-C1 co-expression (iv), indicating its resistance to BoNT-C1, while CSYS;S14A (i) was located in the cytosol upon BoNT-C1 co-expression (ii). Scale bar: 5 mm. *Lower panel:* normalized fluorescence intensity profiles of the above cells indicating PM or cytosolic expression. The fluorescence profiles were determined from line scans (red lines, upper panel) taken from the outside to the middle of each cell. (**B)** Representative amperometric recordings of catecholamine release from cells transfected CSYS (R) or with CSYS;S14A (R), before and after hK depolarization. **(C)** Average number of spikes (**a**) and total charge release (**b**) for spontaneous (before hK) and evoked (after hK) release from PC12 cells co-transfected with BoNT-C and either CSYS (R) or CSYS;S14A (R) (t(14) = 2.16, p = 0.048 (**a**) and t(14) = 2.84, p = 0.013 (**b**), *paired t test*). **(D)** Average number of spikes for spontaneous release from PC12 cells co-transfected with BoNT-C and either CSYS-5RK/A (R) or CSYS-5RK/A;S14A (R) (n= 11 and 8, respectively, P = 0.044, *Mann-Whitney Rank Sum Test*)

Together, the amperometry analysis (Fig. 7) and the FRET analysis of the TeTx effect (Fig. 6D), indicate that S14 phosphorylation is required for Ca^2+^-regulated vesicle exocytosis.

## Discussion

This study demonstrates the importance of CK2 phosphorylation of Syx at S14 in the regulation of a depolarization-induced Ca^2+^-dependent opening of Syx, and in Ca^2+^-triggered release. The role of S14 phosphorylation is highlighted by three major findings obtained using the Syx-based FRET probe, CSYS, previously shown to support Ca^2+^-triggered release (Greitzer-Antes et al., 2013; Singer-Lahat et al., 2018). The first finding was obtained by dynamic FRET analysis of the effects of phospho-null and phospho-mimetic CSYS mutations, as well as the effect of impaired *in vivo S14* phosphorylation, either by disruption of the CK2 recognition site or inhibition of the catalytic site. These experiments establish that S14 phosphorylation is responsible for the previously described (Greitzer-Antes et al., 2013) requirement of the juxtamembrane 5RK stretch of Syx for CDO. The second finding, obtained by dynamic FRET analysis of the effect of the phospho-null mutation in the presence of TeTx LC, which cleaves the vesicular SNARE Syb, reveals that S14 phosphorylation causes CDO to occur within a vesicular context. The third finding, obtained by amperometric analysis of the effects of the phospho-null mutation on secretion phenotypes, shows that S14 phosphorylation is required for release, spontaneous and Ca^2+^-triggered. These findings demonstrate that S14 phosphorylation is crucial for Syx functioning within a vesicular context under the control of 5RK to support Ca^2+^-dependent release. Together with our previous results, which assigned a dual role for 5RK in clamping spontaneous release and stimulating Ca^2+^-triggered release (Singer-Lahat et al., 2018), these findings provide a basis for a mechanistic model (described below) that describes interrelated roles for S14 phosphorylation and 5RK in the functioning of Syx in Ca^2+^-dependent exocytosis.

It should be noted that, although several proteins involved in Ca^2+^-dependent exocytosis, including Syt-1, Syx, VAMP-2, and CAPS, are *in vitro* substrates of CK2, (Bennett et al., 1993; Davletov and Sudhof, 1993; Nielander et al., 1995; Nojiri et al., 2009), only the activity of CAPS has so far been shown to be dependent on CK2 phosphorylation *in vivo*. This study is the first report to directly link *in vivo* CK2 phosphorylation and the activity of Syx as important for exocytosis.

### Model describing the interrelated roles of CK2-induced S14 phosphorylation and 5RK in the functioning of Syx in Ca^2+^-triggered releas

Integrating our FRET and amperometric results, previous (Greitzer-Antes et al., 2013; Singer-Lahat et al., 2018) and current, together with evidence from the recent literature, has allowed us to formulate a number of postulations that provide the basis for a model of Syx functioning in exocytosis (Fig. 8).

**Figure 8.**
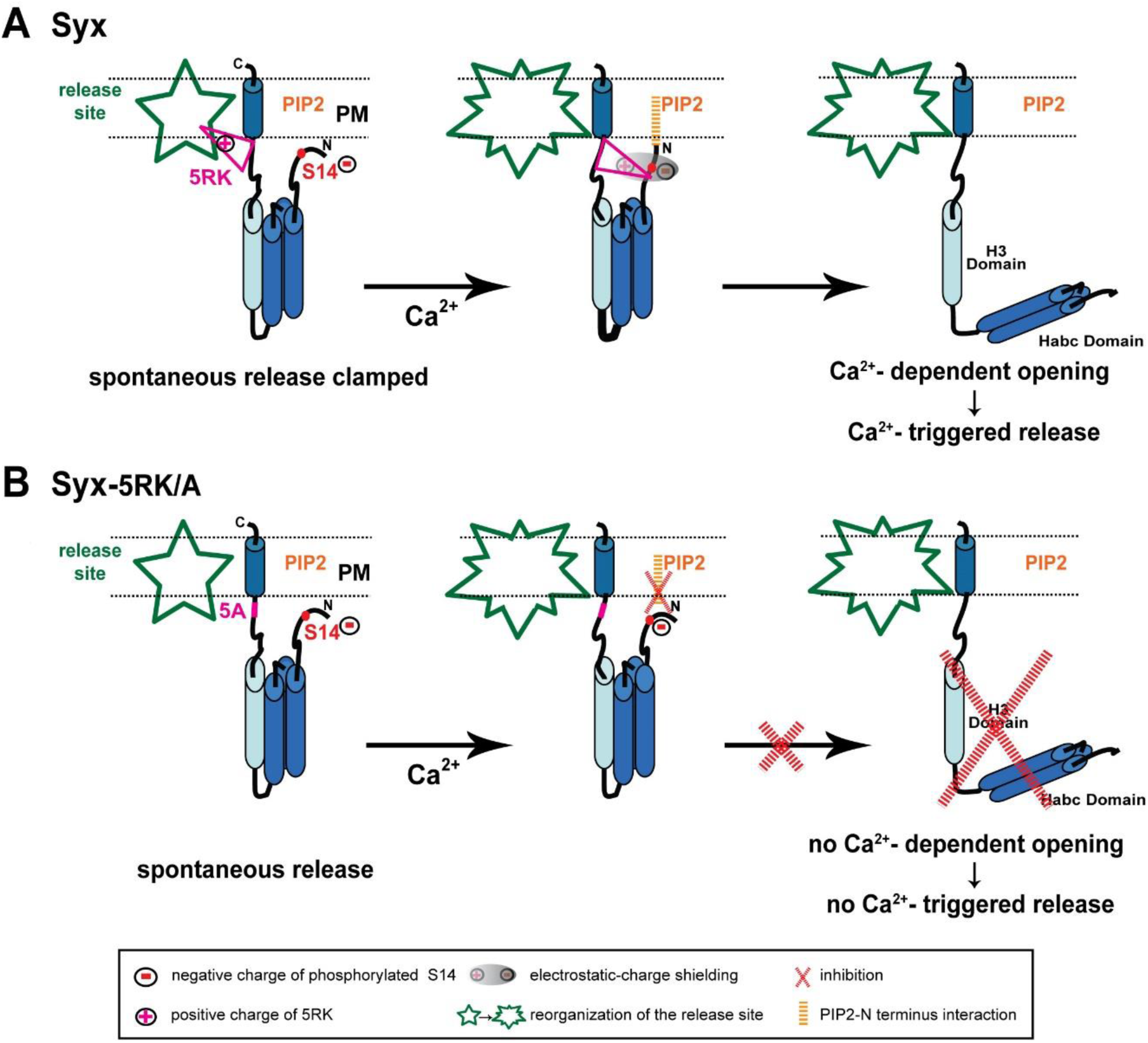
Model of the interrelated roles of 5RK and S14 phosphorylation in Syx functioning during exocytosis. **(A)** *Left panel*, Syx (blue cylinders) maintains the closed conformation at release competent sites (green; release site) due to S14 phosphorylation by CK2. Prior to Ca^2+^ entry, 5RK (pink triangle) interacts with certain lipids and /or fusogenic proteins that enable 5RK to act as a fusion clamp that inhibits spontaneous release. Under these conditions, PIP2 is not likely to interact with Syx (possibly with the N terminus), because of hindrance by the negative charge of the phosphate group of phosphorylated S14. *Middle and right panels*, Sequential reactions in response to depolarization-induced intracellular Ca^2+^ elevation start with reorganization of the release site and an intramolecular rearrangement of Syx that draws the positive charges of 5RK and the negative charges of phosphorylated S14 into close proximity, resulting in electrostatic-charge shielding (grey shade) of the phosphate group (*middle*). The consequent formation of the PIP2–Syx interaction (orange) enables Syx to undergo CDO and to promote Ca^2+^-triggered release (*right*). **(B)** *Left panel*, the S14-phosphorylated 5RK/A mutant lacks the 5RK clamp and exhibits enhanced spontaneous release. *Middle and right panels*, In response to depolarization-induced intracellular Ca^2+^ elevation there is no charge shielding of the phosphate group so that PIP2 cannot interact with Syx, consequently Syx cannot undergo CDO and promote Ca^2+^-triggered release. (PM; plasma membrane).

First, in spite of the documented importance of the electrostatic interactions between PIP2 and 5RK in vesicle exocytosis (Introduction; reviewed in (Martin, 2015)), the interaction between PIP2 and Syx identified in our study proved to be unrelated to 5RK. We demonstrated that, while PIP2 is essential for CDO in both the S14 phosphorylated (Singer-Lahat et al., 2018) and the non-phosphorylated (Fig. 6) states of CSYS, 5RK is essential for CDO only in the phosphorylated state ((Singer-Lahat et al., 2018) and Figs. 2-5). This means that the roles of 5RK and PIP2 in CDO are distinct and do not involve the commonly assumed PIP2-5RK electrostatic interaction. Indeed, molecular dynamics simulations suggested that sites in Syx that are critical to its interaction with PIP2 are not confined to 5RK; in the closed conformation of Syx, the N terminus presumably lies in a juxtamembrane position that enables it to interact with PIP2 (Khelashvili et al., 2012). Taken together, *the first postulation is that an inherent PIP2-Syx interaction, possibly involving the N terminus of Syx, is absolutely required for CDO, regardless of CK2 phosphorylation*.

Second, we note that CK2 phosphorylation has been reported to play a significant role in neuronal excitability in a location-specific manner. Specifically, CK2 phosphorylation has been shown to regulate the accumulation of voltage gated Na^+^ (Brachet et al., 2010; Hien et al., 2014) and K^+^ (Adelman et al., 2012; Allen et al., 2007) channels at the axon initial segment, with great impact on axonal excitability, and to affect NMDA receptors in a receptor-location-specific manner to modulate long term potentiation (Kimura and Matsuki, 2008). Notably, location-specificity was reported also for S14 phosphorylated Syx, shown to localize to discrete domains along a subset of axons (Foletti et al., 2000). Here we show that, in contrast to CSYS (being endogenously S14 phosphorylated; Fig 1Bb), the phospho-null CSYS mutant undergoes CDO that does not involve the vesicular SNARE, Syb2 (Fig. 6D) and does not support Ca^2+^-triggered release (Fig. 7A-C). Furthermore, in contrast to CSYS-5RK/A (being endogenously S14 phosphorylated; Fig. 5Ab), the phospho-null CSYS-5RK/A mutant does not cause prominent spontaneous release (Fig. 7D). These observations allow us to associate the determined location-specificity of S14 phosphorylated Syx with release-competent sites, where Syx acts to support vesicle release events, spontaneous and Ca^2+^-triggered. Thus, *the second postulation is that S14 phosphorylated Syx is specific to release-competent sites. In these sites Syx can be engaged in ternary SNARE complex assembly and support release*.

Third, the previously described secretion phenotype of CSYS-5RK/A, with enhanced spontaneous release, suggested that 5RK serves as a fusion clamp (Singer-Lahat et al., 2018). Thus, *the third postulation is that 5RK acts to clamp spontaneous release at release-competent sites*, presumably via its well documented interactions with lipids and/or fusogenic proteins (*Introduction*).

Fourth, since CSYS-5RK/A cannot undergo CDO (Greitzer-Antes et al., 2013), unless phosphorylation of its S14 is impaired (Figs. 2C and 5B), *the fourth postulation is that S14 phosphorylation inhibits CDO in the absence of intact 5RK; 5RK relieves the inhibition*, as exemplified in CSYS (with intact 5RK), which undergoes CDO while being phosphorylated. We hypothesize that the mechanism underlying the inhibition is the formation of an electrostatic repulsion between the negatively charged phosphate group of phosphorylated S14 and PIP2, interfering with the N-terminal-PIP2 interaction that is essential for the ability of Syx to undergo CDO (first postulation). This inhibition can be relieved upon masking of the negative charge of the phosphate moiety by the positive charge of 5RK (both charges presumably lie in juxtamembrane positions). The charge masking may require Ca^2+^-induced structural reorganization(s) of Syx, possibly with respect to the phospholipid environment (for example see: (Milovanovic et al., 2016)), that bring 5RK and the phosphate moiety into close proximity with consequent electrostatic-charge shielding.

Fifth, since Ca^2^-triggered release, similarly to CDO, is induced only in the presence of 5RK, our *fifth postulation is that the relief by 5RK of S14 phosphorylation-induced inhibition of CDO (fourth postulation) underlies Ca*^*2+*^*-triggered release*.

Based on the above postulations, we present the following model of interrelated roles of 5RK and S14 phosphorylation in Syx functioning during exocytosis (Fig. 8A). S14 phosphorylated Syx, in contrast to its non-phosphorylated form (not shown in the model; see below), resides in release-competent sites. Prior to Ca^2+^ entry, Syx assumes its closed conformation, with 5RK serving as a fusion clamp that inhibits spontaneous release, possibly via interactions with phospholipids and/or fusogenic proteins (Fig 8A, *left*). In response to a depolarization-induced intracellular elevation of Ca^2+^, an intramolecular rearrangement of Syx, possibly associated with the reorganization of the release-competent site, draws 5RK and phosphorylated S14 into close proximity. The resulting electrostatic-charge shielding of the phosphate group by 5RK permits an otherwise inhibited interaction of the N terminus of Syx with PIP2, which is essential for CDO (Fig 8A, *middle*). Subsequently, Syx undergoes CDO and promotes Ca^2+^-triggered release (Fig. 8A *right*). In contrast, the 5RK/A mutation of Syx (Fig. 8B) enhances spontaneous release, due to lack of the 5RK fusion clamp (Fig. 8B, *left*), while Ca^2+^-triggered release is inhibited, because the PIP2-Syx interaction, which enables CDO, cannot occur while the phosphate charge is not masked by 5RK (Fig. 8B, *middle & right*).

In conclusion, we suggest that CK2-induced S14 phosphorylation of Syx is essential for vesicle release as it localizes Syx to a release-relevant setting in which it acts in concert with 5RK to make release dependent on depolarization-induced intracellular Ca^2+^ elevation.

According to our paradigm, non-phosphorylated Syx (not shown in the model) is miss-located to a release-irrelevant environment. There, it undergoes CDO that does not include Syb2 (Fig. 6D), likely reflecting the formation of Ca^2+^-induced non-productive SNARE intermediate(s) (e.g. 2:1 Syx/SNAP-25 binary complexes; (Hernandez et al., 2012; Weninger et al., 2008)), and does not support vesicle fusion (Fig. 7). Notably, in the absence of S14 phosphorylation, there is no electrostatic repulsion between the N terminus and PIP2 so that CDO can occur regardless of whether intact 5RK is present (Fig. 2). Namely, CDO regulation by 5RK is confined to release-relevant sites.

## Materials and methods

### Plasmid construction

Double-labeled Syx (CSYS) and CSYS-5RK/A cDNA were generated as described in (Greitzer-Antes et al., 2013). CSYS;S14A was generated by introducing a point mutation, S14A, in the N-peptide domain of CSYS. CSYS;5D/A was generated by introducing 5 point mutations, D15,16,17,18,19A, in the N-peptide domain of CSYS. The BoNT-C1-resistant mutation (CSYS(R)) was generated by introducing one point mutation, K253I, in the BoNT-C1 recognizing sequence (Binda et al., 2008). For PC12 transfection, the constructs were cloned into pcDNA3 vector using EcoRI and XbaI restriction sites. The PLCη2 construct was kindly provided by T.F.J. Martin (University of Wisconsin). BoNT-C1α51 was subcloned into an mRFP pcDNA3 vector containing IRES (internal ribosome entry site).

### PC12 cells preparation and transfection for FRET experiments

PC12 cells were maintained at 37°C/5% CO_2_ in Dulbecco’s Modified Eagle’s Medium (DMEM) with high glucose (Sigma-Aldrich) supplemented with 10% Bovine serum, 5% L-glutamine, 100 U/ml penicillin, and 0.1 mg/ml streptomycin. For imaging, cells were replated to ∼60% confluence on poly-L-Lysine (Sigma-Aldrich) coated 35 mm glass bottom culture dishes, and allowed to adhere overnight. Cells were transfected with 1.5 μg of the CSYS mutation probes, using Lipofectamine 2000 (Invitrogen). Imaging experiments were conducted at room temperature, 24 h after transfection. During the experiment, the transfected cells were superfused through a 2 ml bath, with physiological (2.8 mM K^+^) and high K^+^ (105 mM K^+^) solutions as described previously (An and Almers, 2004)

### Dynamic FRET assay

PC12 cells were imaged using a C-Apochromat 40x/1.2 NA water objective and excited with a 405-nm laser every 5 sec for a total of 400 sec. During the sequential imaging, the cells were imaged in a control, 2.8 mM K^+^, solution for 100 sec, before and after 200 sec of 105 mM high K^+^ solution, for a total imaging time of 400 sec. Fluorescent signals were collected with a Zeiss 510META confocal microscope using its “channel mode”. Cells were excited with a 405 nm laser band and the emission was filtered through the main beam splitter HFT405/514/633 nm and further separated by a secondary beam splitter, NFT515 nm. CFP and YFP fluorescence were collected by a 470-500 nm and 505-550 nm band pass filter, respectively, and directed into two separate photomultipliers. Under these settings, the leak of YFP into the CFP recording window is purely optical, and very low (< 1%) (Okamoto et al., 2004), and remains constant regardless of changes in FRET, thus it does not require any corrections. YFP and CFP intensities in a region of interest (ROI) on the cell membrane were calculated, and background fluorescence was quantified from an ROI defined in each image in an area containing no fluorescent cells. The background-subtracted fluorescence intensity at each exposure time point was normalized to the average of the initial measurements in each cell (before the high K^+^ solution was added). The FRET ratio of normalized intensities was denoted as F_YFP_/F_CFP_. Changes in FRET are reflected as changes in the FRET ratio.

### Cell culture for the amperometry experiments

PC12 cells were obtained from the American Type Culture Collection (ATCC, Rockville, MD), and maintained in RPMI-1640 media supplemented with 10% heat-inactivated horse serum and 5% fetal bovine serum (Sigma-Aldrich), in a 7% CO_2_ atmosphere at 37°C. The cells were grown on mouse collagen IV-coated flasks (Becton Dickinson, Bedford, MA) and were subcultured approximately every 7 days. Cells were transfected using Lipofectamine-2000 (Invitrogen).

### Amperometry measurements

Amperometric currents were recorded with a VA-10 amplifier (npi Electronics GmbH, Tamm, Germany). A constant voltage of +700 mV was applied to the 5 μm OD carbon electrode (ALA Scientific, Westbury, NY). The currents were filtered at 1 kHz and digitized at 10 kHz using a Digidata 1322A analog-to-digital converter and the Clampex 9 software package (Axon CNS Molecular Devices, Sunnyvale, CA). Data were analyzed by the customized macro for IGOR PRO software (Mosharov, 2008). Spikes smaller than 5 pA were considered to be noise and were discarded. The bath solution contained (in mM): 150 NaCl, 5 KCl, 1.2 MgCl_2_, 5 glucose, 10 Hepes, 2 CaCl_2_, pH 7.4, and the high K^+^ solution contained (in mM): 100 KCl, 50 CaCl_2_, 0.7 MgCl_2_, Hepes 10, pH 7.4. Similarly to what was shown in chromaffin cells (Shang et al., 2014), we observed reduced spontaneous release in the presence of elevated [Ca^2+^] in PC12 cells (data not shown); therefore, spontaneous events were monitored in physiological [Ca^2+^] (2 mM).

### Immunoprecipitation (IP) and Immunoblotting (IB) in PC12 cells

PC12 cells were homogenized in the lysis buffer: 150 mM NaCl; 50 mM Tris-HCl; 5 mM EDTA; 1 mM EGTA; 10 mM sodium fluoride and 1 mM sodium orthovanadate, containing a cocktail of protease inhibitors (Promega). Following solubilization by 1% Triton X-100, the homogenate was centrifuged (12000xg/25 min/4°C) and the supernatant was incubated with YFP (MBS 833304) antibody overnight. Proteins bound to the beads were eluted by heating at 95° C for 4 min in the SDS-PAGE sample loading buffer. Immunoprecipitated proteins were separated by SDS-PAGE and subjected to Western blotting with antibody against S14 phosphorylated Syx (Abcam; ab63571), using the ECL detection system (Pierce Protein Research Products, Thermo Scientific). In experiments investigating the effect of TBB, cells were incubated with 20 µM of TBB (Merck) for one hour prior to homogenization.

### Immunoprecipitation (IP) and Immunoblotting (IB) in *Xenopus* oocytes

de-folliculated *Xenopus* oocytes were metabolically labeled 4 h after mRNA injection, by incubation in NDE solution containing 0.3 mCi/ml of ^35^ [S] methionine/cysteine for 2 days. After this time, 6 to 8 oocytes were homogenized in 1 ml of medium composed of 20 mM Tris (pH 7.4), 5 mM EDTA, 5 mM EGTA, and 100 mM NaCl [containing protease inhibitor cocktail (Roche)]. Debris was removed by centrifugation for 10 min at 4°C. After solubilization by 1% Triton X-100, the homogenate was centrifuged (12000xg/15 min/4°C) and the supernatants were incubated with anti-Syx antibody (Alomone labs) overnight at 4°C with gentle rotation. Immunoprecipitated proteins were separated by SDS-PAGE and subjected to Western blotting with antibody against Syx phospho-S14 (Abcam; ab63571). Digitized scans were derived by PhosphorImager (Molecular Dynamics, Eugene, OR) and relative intensities were quantitated by ImageQuant.

### Experimental design and Statistical Analysis

Data was summarized as means ± S.E.M. The number of samples (n) indicates the number of cells per group. Statistical analysis was performed in either SPSS v24 (IBM) or SigmaPlot v11. One-way analysis of variance (ANOVA) for repeated measures was used to analyze time course FRET (dynamic FRET) experiments. For amperometry experiments where events were monitored before and after stimulation in each cell, paired *t*-test analysis was used. For 2 independent groups of events with non-Gaussian distributions, we used the Mann-Whitney Rank Sum Test. Multiple groups were compared by ANOVA followed by post-hoc Tukey’s test. Asterisks in the figures indicate statistically significant differences as follows: * p < 0.05; ** p < 0.001.

